# Specter: linear deconvolution as a new paradigm for targeted analysis of data-independent acquisition mass spectrometry proteomics

**DOI:** 10.1101/152744

**Authors:** Ryan Peckner, Samuel A Myers, Jarrett D Egertson, Richard S Johnson, Jennifer G. Abelin, Steven A Carr, Michael J MacCoss, Jacob D Jaffe

**Author notes:** Correspondence should be addressed to J.D.J.

## Abstract

Mass spectrometry with data-independent acquisition (DIA) has emerged as a promising method to greatly improve the comprehensiveness and reproducibility of targeted and discovery proteomics, in theory systematically measuring all peptide precursors within a biological sample. Despite the technical maturity of DIA, the analytical challenges involved in discriminating between peptides with similar sequences in convoluted spectra have limited its applicability in important cases, such as the detection of single-nucleotide polymorphisms and alternative site localizations in phosphoproteomics data. We have developed Specter, an open-source software tool that uses linear algebra to deconvolute DIA mixture spectra directly in terms of a spectral library, circumventing the problems associated with typical fragment correlation-based approaches. We validate the sensitivity of Specter and its performance relative to other methods by means of several complex datasets, and show that Specter is able to successfully analyze cases involving highly similar peptides that are typically challenging for DIA analysis methods.

## 1 Introduction

Mass spectrometry with data-dependent acquisition (DDA) is the method of choice for large-scale discovery proteomics, enabling the rapid measurement of the abundances of thousands of proteins in a sample without any prior knowledge of its contents. While it is a powerful technique for the identification of high-abundance proteins in individual samples, DDA faces a fundamental limit in terms of reproducibility and comprehensiveness due to the stochastic nature of its data gathering process^1^. This inhibits the consistent detection of proteins across samples and undermines efforts to identify particular proteins as biomarkers of disease or measure their differential abundance between multiple conditions of interest. As a result, targeted strategies such as parallel reaction monitoring (PRM) or selected reaction monitoring (SRM) have been developed in order to guarantee the measurement of low-abundance analytes or the observation of pre-specified targets across multiple samples^2^, but this gain in specificity comes at the cost of vastly limiting the range of observable precursors.

Data-independent acquisition (DIA) is a relatively new approach that combines the reproducibility of SRM with the breadth of DDA by simultaneously fragmenting all precursors whose mass-to-charge (*m/z*) ratios fall into one of a small number of wide windows that traverse the entire *m/z* range. This results in convoluted MS2 spectra whose fragment ion intensities may be comprised of contributions from multiple peptide precursors and which are far more complex to analyze than their DDA counterparts.

The new challenges posed by DIA demand specialized software tools for downstream analysis^3-7^. Most of these are targeted methods that require a user-provided spectral library defining the search space of peptides (and, in turn, proteins) that can be identified and quantified in the acquired data. These targeted tools are for the most part derived heuristically from corresponding methods for DDA and PRM analysis, in which library members are scored against acquired MS2 spectra based on characteristics such as normalized dot product, fragment ion correlation, and chromatographic peak shape. Although these scores typically penalize assignments to library spectra whose annotated *b* or *y* fragment ions are judged to exhibit interferences, these methods do not rigorously account for the confounding effects of precursor cofragmentation, resulting in a limited ability to distinguish precursors with shared spectral features. Alternatively, untargeted methods^4,5,8^ aim to deconvolute the data directly without the use of spectral libraries based on the grouping of fragment ions with correlated elution profiles. This analysis takes into account both MS1 and MS2 intensity information in order to score correlated peak groups but implicitly discards fragments with significant interferences due to their poor correlation with a precursor’s elution profile. Although a promising route for the discovery of previously unobserved analytes, the fact that this approach uses no prior information as provided by a spectral library may cause it to suffer from a high false negative rate in complex samples^9^ and makes it more susceptible to missing data than targeted methods when attempting to quantify analytes across multiple conditions.

Here we describe Specter, an algorithm for the identification and quantification of spectral library members within DIA data. It is based on recognizing and formalizing the fundamental distinction between DIA and DDA, namely the cofragmentation of potentially large numbers of precursors, some of which may share fragment ion *m/z* ratios. Specter is based on a principled mathematical formulation of the cofragmentation problem, which is then solved by means of linear algebra. As opposed to the usual approach involving detection of correlated chromatographic profiles of selected precursor fragment ions, spectral deconvolution takes place purely at the MS2 level and involves the entire sequence of *m/z* coordinates and relative intensities of library spectra peaks. This allows for the direct calculation of extracted ion chromatograms of fractional contributions of library precursors to MS2 spectra, which can then be visualized and analyzed using traditional chromatographic approaches^10^.

This approach is neither spectrum-centric (it does not match acquired MS2 spectra to those in a database) nor peptide-centric (it does not assess the evidence for individual library members within the acquired data), in contrast to all other existing methods. Rather, it is “combination-centric”, in that it identifies and quantifies the single *combination* of library spectra that best explains an acquired MS2 spectrum. This approach removes the need to reduce spectra to curated fragment ions, carries an intrinsically low false discovery rate, and allows us to distinguish precursors with highly similar library spectra such as those originating from single-nucleotide polymorphisms or positional isomers in phosphoproteomics data. Moreover, the linear algebraic framework establishes a meaningful notion of the quantification of a precursor within a single DIA MS2 spectrum independently from chromatographic information. Specter is able to analyze DIA-type data from any instrument vendor and acquisition scheme (e.g. SWATH or MS^*E*^), and requires only three inputs: an experimental data file in centroided mzML format, a spectral library in Bibliospec’s blib format^11^, and a user-specified mass accuracy parameter (either as an absolute *m/z* value or a parts-per-million tolerance). Retention time information in the library is optional, and retention time normalization is not required, though it might improve the speed and quality of the results. Specter is built on the open-source distributed computing framework Apache Spark and is available as an open-source software tool at https://github.com/rpeckner-broad/Specter.

## 2 Results

### 2.1 Specter is based on an algebraic approach to deconvolving mixed spectra

Specter is based on the hypothesis that every MS2 spectrum *S* acquired in the course of a DIA run is a *linear combination* of the spectra of the precursors that are cofragmented to acquire it (disregarding the effects of biochemical noise, instrument error, and experimental variability in a peptide’s fragmentation pattern). This is because the total number of ions with a particular *m/z* ratio in *S* is, in ideal terms, simply the sum of the number of ions with that *m/z* contributed by each of the constituent precursors of S. Furthermore, the number of ions with a certain *m/z* that are produced by any one of these cofragmented precursors is entirely determined by the precursor’s fragmentation pattern (its pure spectrum) and abundance at the time *S* is acquired.

The recognition that mixed mass spectra are linear combinations of pure ones has long been an established principle in gas chromatography-mass spectrometry^12,13^ and, more recently, metabolomics^14,15^. However, so-called matrix methods for spectral deconvolution have for the most part not taken shape in usable software for GC-MS applications^13^, and to the best of our knowledge Specter is the first software tool to apply linear deconvolution to mass spectrometry proteomics data.

The linear combination principle is illustrated by Figure 1 and takes the form of the matrix equation

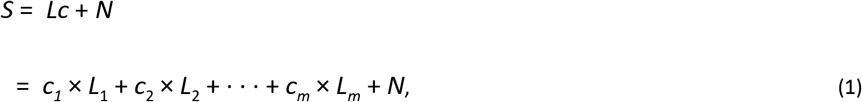

where *S* is a vector containing the intensities of the peaks of the DIA MS2 spectrum at their respective *m/z* values; *L* is a matrix whose columns are the library spectra *L*_1_ *L*_2_, …, *L_m_* under consideration, normalized so that their total ion intensities are all equal to one; *N* is a vector of noise whose components are unknown; and the vector *c = (c*_1_ *c*_2_, …, *c_m_*), which Specter calculates by nonnegative linear regression (Methods §4.4), describes the quantitative contribution of each library spectrum to the mixed spectrum *S* ( see §2.1.1 below).

Equation (1) formalizes the fundamental difference between DIA and DDA, namely the cofragmentation of multiple precursors, by treating precursor cofragmentation mathematically as the addition of library spectra (Fig. 1). The linear algebra framework allows us to calculate the quantitative contribution of each library precursor (constituting a column of the matrix L) to each MS2 spectrum in terms of the coefficients *c_i_*. These algebraic coefficients are calculated for each MS2 spectrum and then analyzed further to determine the final identifications and quantifications of library members.

**Figure 1:**
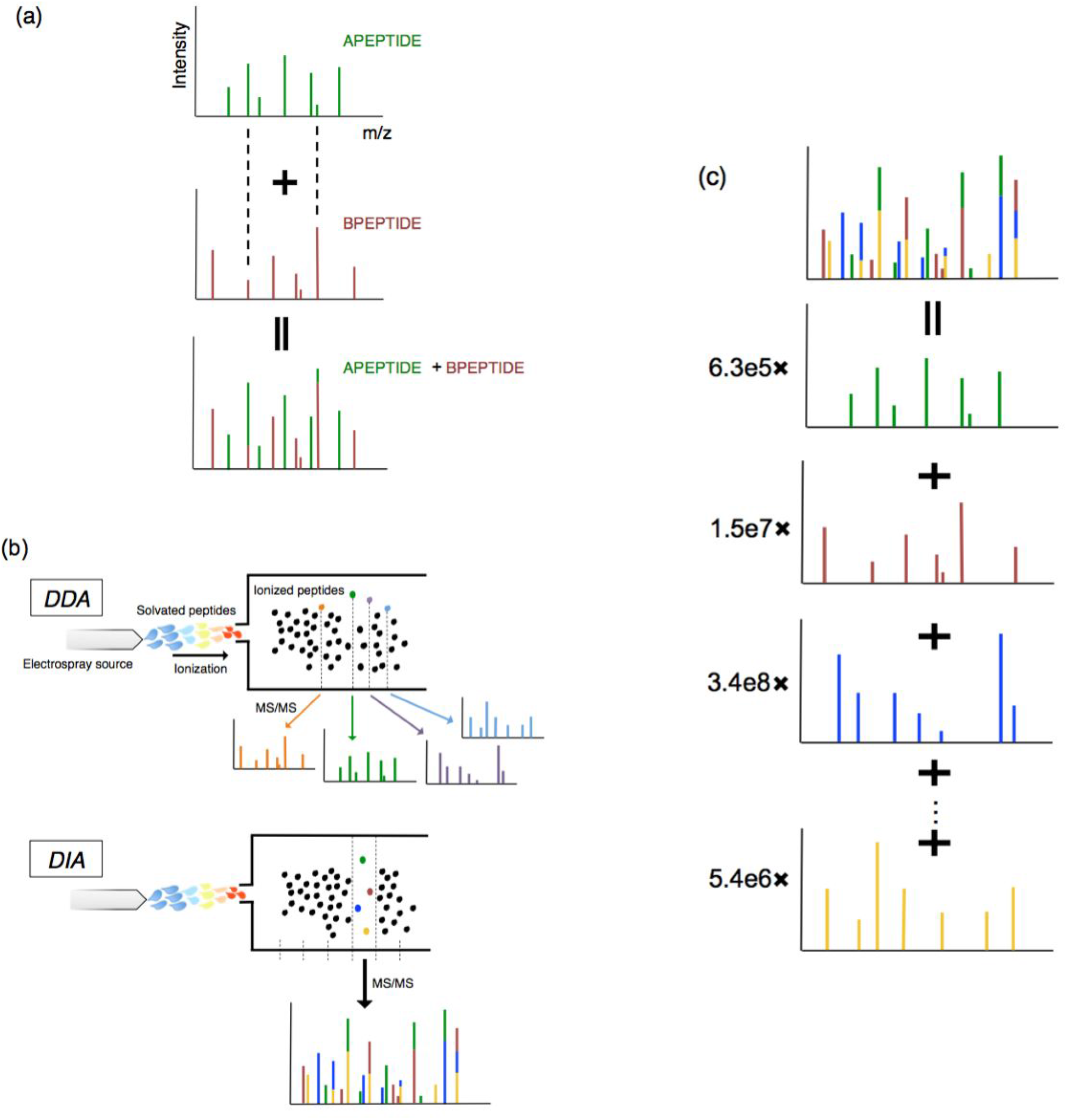
Specter uses linear algebra to formally deconvolute MS2 spectra derived from cofragmented precursors. (a) Mixed spectra are linear combinations of pure spectra. Pure spectra of two hypothetical precursors are shown. Their cofragmentation results in addition of their pure spectra to obtain a mixed spectrum containing fragment ion intensities from both precursors. Fragments with identical *m/z* ratios in the two pure spectra, whose positions are indicated by the dashed lines, lead to peaks in the mixed spectrum whose intensities are composed of contributions from both precursors. (b) In DDA, precursors are selected in decreasing order of abundance and fragmented separately to form MS2 spectra that are typically composed of a single precursor. In DIA, groups of precursors whose *m/z* ratios fall into the same wide window are fragmented simultaneously to form mixed MS2 spectra. (c) Specter finds the quantitative combination of library spectra that most closely matches the acquired DIA spectrum by linearly deconvolving the mixed spectrum into pure components from the library. The coefficient of each library spectrum is the total ion intensity of the corresponding precursor within the acquired spectrum.

#### 2.1.1 Specter coefficients are readily interpreted as total ion intensities

Protein quantification is critical for the inference of regulatory function and discovery of disease biomarkers. The total ion intensity of a precursor in an MS2 spectrum, meaning the sum of the absolute intensities of all fragment ions produced by that precursor in the spectrum, is the basic unit of quantification in label-free DIA mass spectrometry. This is in contrast to the MS1-based intensity typically employed in DDA, where most MS2 spectra are comprised of a single precursor (or, occasionally, a small number thereof). Total ion intensity determined at the MS2 level is a more sensitive mode of quantification for DIA spectra comprised of multiple cofragmented precursors, as some of these may be of too low an abundance to be reliably quantified at the MS1 stage.

The series of Specter coefficients associated to a given library member has a natural interpretation as a calculated total ion chromatogram (Methods §4.5). We measure the overall similarity of this calculated chromatogram to an ideal Gaussian-like elution profile by scoring its height, variance, skewness and kurtosis (Methods §4.5). Scores of this type are commonly used for quality assessment of total ion chromatograms^10^, and we show below in §2.2.1 that they accurately differentiate the Specter chromatograms of false positive decoy identifications from those of true positive precursors.

### 2.2 Specter is as accurate as targeted manual analysis in terms of both identification and quantification

We used Specter to analyze DIA data generated as part of a study to compare the quantitative performance of different DIA methods^16^. These data were obtained by spiking a commercially-available digest of five equimolar bovine proteins into an *S. cerevisiae* lysate digest at ten levels ranging from 0 amol to 30 fmol per injection, and analyzing each spike-in sample in triplicate on a Q-Exactive Orbitrap HF using DIA (Methods §4.8). Specter was applied to each resulting data file using a spectral library consisting of 887 yeast and 72 bovine precursors that was acquired from a DDA run of the 30 fmol spike-in sample. Independently, a targeted manual analysis of the unprocessed data for the bovine peptides was performed in Skyline using the same library.

Specter accurately calculates the total ion intensities of each of the bovine peptides across the spike-in concentrations and replicates (Fig. 2). The elution profiles calculated by Specter agree extremely well with expert annotation in terms of both total ion intensities and the retention times at which they are identified (Figure 2(a) and Supplementary Fig. 1). Furthermore, the quantifications derived by calculating the area under the calculated Specter chromatogram for each precursor (Methods §4.6) exhibit the expected linear increase with spike-in concentration (Figure 2(b)). They also agree with manual expert quantification, with the Pearson correlation for all fifty-five peptides from all of the bovine proteins over all spike-ins over the limit of detection of 300 amol being 0.93 (Figure 2(c)). While manual quantifications are determined from the extracted ion chromatograms of only pre-selected fragment ions, quantification by Specter incorporates every peak present in its library spectrum. Thus, some differences between the quantifications are expected. Furthermore, by analyzing the same data with three different libraries, we found that the coefficients calculated by Specter are robust with respect to spectral library noise and incompleteness (Supplementary Figure 1).

**Figure 2:**
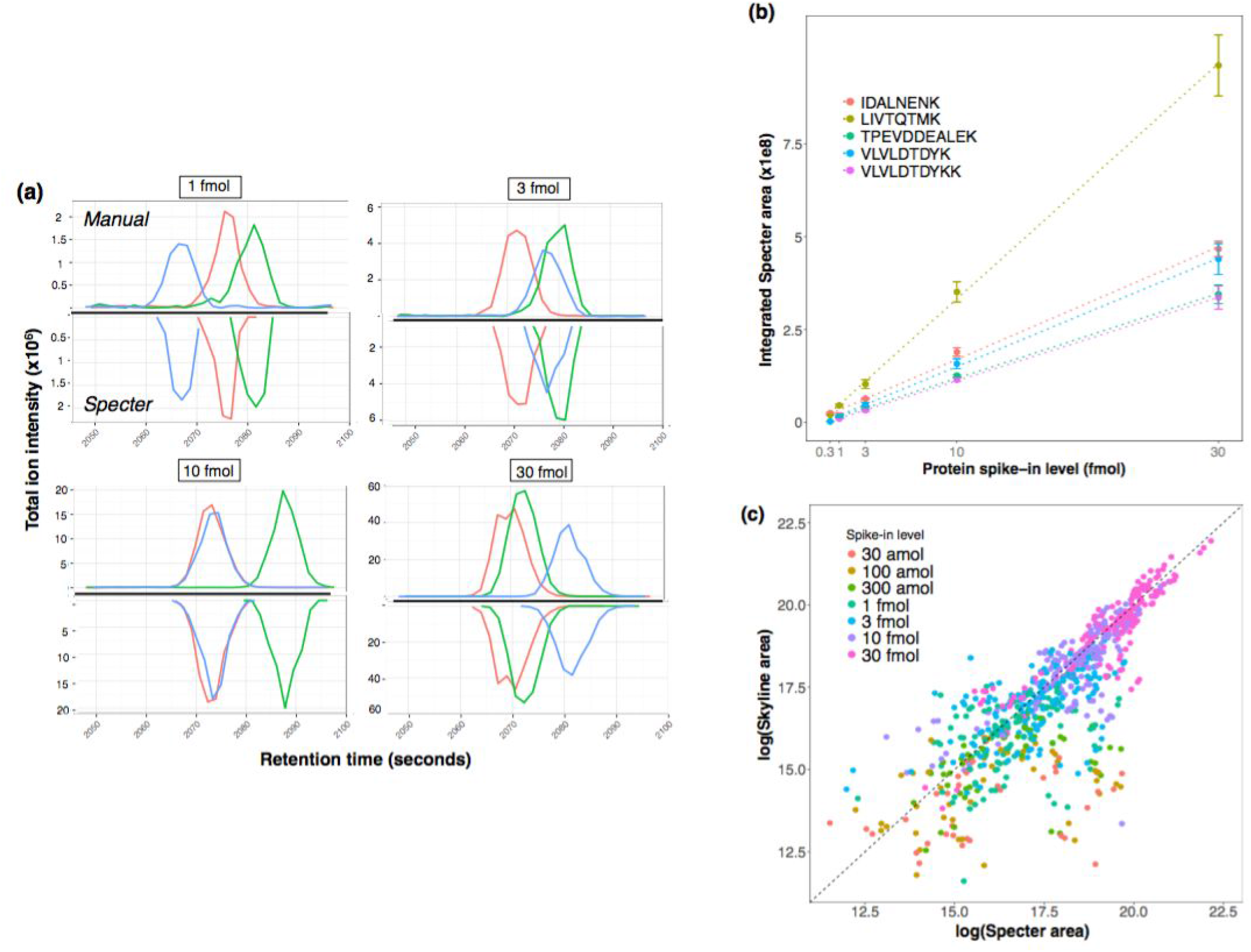
Total ion chromatograms calculated by Specter are as accurate as those from manual targeted analysis of DIA data for both identification and quantification. Bovine proteins were spiked into yeast lysate over an increasing range of concentrations, and each spike-in sample was analyzed in triplicate in DIA mode. (a) Total ion chromatograms determined by manual quantification in Skyline for the peptide VLVLDTDYKK from the bovine protein *β*-lactoglobulin, mirrored by those calculated by Specter, for four spike-in levels. Each colored line within each panel describes the total ion chromatogram for this precursor within a single replicate run. Note the increasing y-axis scales. (b) Means of quantifications by Specter (area under the Specter chromatogram) over replicates for the five identified peptides from bovine *β*-lactoglobulin over the range of spike-in volumes, together with linear fits (dotted lines). Error bars indicate standard error across replicates. (c) Quantifications by Specter vs. manual for all fifty-five identified peptides from all bovine proteins in all replicates.

#### 2.2.1 The false discovery rate of Specter is intrinsically low

We adopted the common target/decoy approach^17,18^ to assess the false discovery rate of Specter. We expect that, when decoys are added to a spectral library, at most a very small percentage of precursors initially identified by Specter should be decoys even before cutoffs on chromatographic scores are introduced. “Initially identified” means that the precursor’s sequence of Specter coefficients exhibits a peak consisting of at least five coefficients greater than 1 in consecutive MS2 spectra (Methods §4.5)

Instead of using computationally generated decoy spectra, we augmented the focused spectral library with data from an *E. coli* spectral library, although no E. coli proteins were present in the sample. We then examined the identifications and quantifications calculated by Specter for *E. coli* peptides (composing 98% of the combined library). Our expectation was that this large, experimentally acquired decoy library would pose a more substantial challenge than a smaller library consisting of artificial spectra, both due to its size relative to the “on-target” library and because its members have general spectral features that may be lacking in synthetic decoys (such as isotope distributions and fragment ion traces not of *b* or *y* type). Of the 959 yeast/bovine library precursors, 757 were identified by Specter in all three replicate DIA runs of the 30 fmol spike-in sample (78% of the yeast/bovine library), while 381 of the 48,131 *E. coli* precursors were identified in all three replicates (0.8% of the *E. coli* library), yielding a 1% false discovery rate (Methods §4.8.1). We have also implemented a target-decoy strategy based on constructing synthetic spectra when Specter is applied in general (Methods §4.7 and Supplementary Figure 3).

A summary comparison of the identifications and quantifications calculated by Specter for the yeast and bovine library precursors versus the *E. coli* decoys is shown in Figure 3. Once the Specter coefficients for all precursors and MS2 spectra have been calculated according to Equation (1), a set of four scores for each precursor is calculated based on the shape and height of the total ion chromatogram formed by the time series of that precursor’s Specter coefficients (§2.1.1, Methods §4.5, and Supplementary Note 1.3). These are then combined into a single score by means of linear discriminant analysis^19^. Based on these linear discriminant scores, Specter clearly distinguishes the yeast/bovine library precursors from those in the *E. coli* library (Fig. 3(b),(c)). While 381 false positive *E. coli* identifications have scores above the threshold of 2 for identification with a 1% FDR, they are lower overall than those of the 757 true positives (Fig. 3(c)), so that a more stringent FDR could be applied without a dramatic loss of true positive identifications.

**Figure 3:**
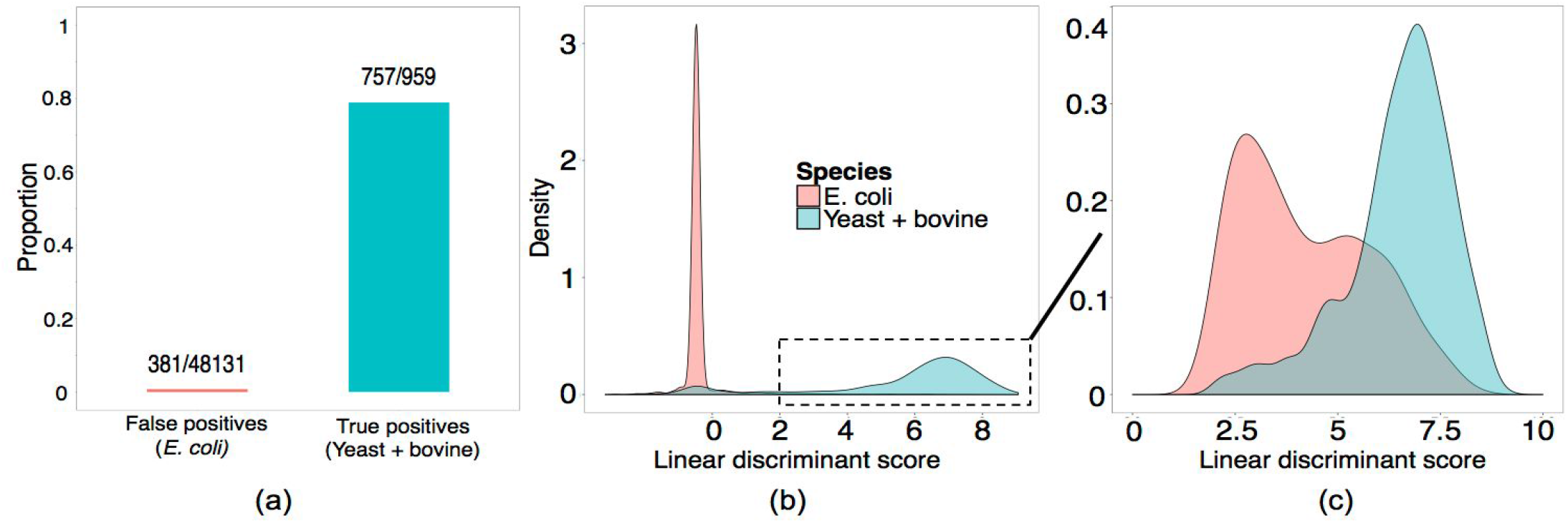
The false discovery rate of Specter is inherently below 1%. (a) Specter identifies a far higher proportion of true positive yeast/bovine library members than false positive *E. coli* library members (79% vs 0.8%). (b) Distributions of linear discriminant scores based on Specter coefficients of all precursors from both libraries. (c) Distributions of linear discriminant scores for only identified precursors (linear discriminant score > 2).

### 2.3 Specter is able to distinguish precursors with highly similar spectra

Spectral libraries may often contain spectra that share a significant number of peaks. Biological processes, such as non-synonymous single-nucleotide polymorphisms (SNPs) in coding regions or alternative localizations for post-translational modifications (PTMs), can result in peptides whose spectra contain a paucity of discriminating fragment ions. Ambiguous shared features are typically deemed ‘interferences’ and excluded from consideration in analysis of DIA data. Such a strategy runs the risk of limiting biological insight, as differential expression of certain SNPs or PTMs may lie at the heart of some disease phenotypes^20^. We aimed to test Specter’s ability to distinguish between extremely similar library spectra with large numbers of shared fragments and to compare this analysis to results obtained using the normalized dot product, the tool used by most other targeted DIA analysis methods to quantify spectral similarity.

We designed an experiment to perform DIA analysis of groups of synthetic peptides whose sequences differ only in single amino acids or by the transposition of a pair of adjacent amino acids. We used three families of synthetic peptides, each consisting of precursors whose spectra are highly similar and whose *m/z* ratios (in charge state +2) fall into the same isolation windows for DIA (Fig. 4 and Supplementary Table B). We then prepared a series of three mixtures (Methods §4.9), in each of which a random set of members of each family was chosen to be spiked into both an *E. coli* lysate digest and a neat background. Each spike-in was then acquired via DIA mass spectrometry, for a total of twelve DIA runs (duplicate runs of both the *E. coli* and neat background samples for three distinct spike-in mixtures). Analysis was performed by Specter using a spectral library consisting of 48,131 E. coli precursors together with the spectra of the synthetic peptides (Methods §4.9).

**Figure 4:**
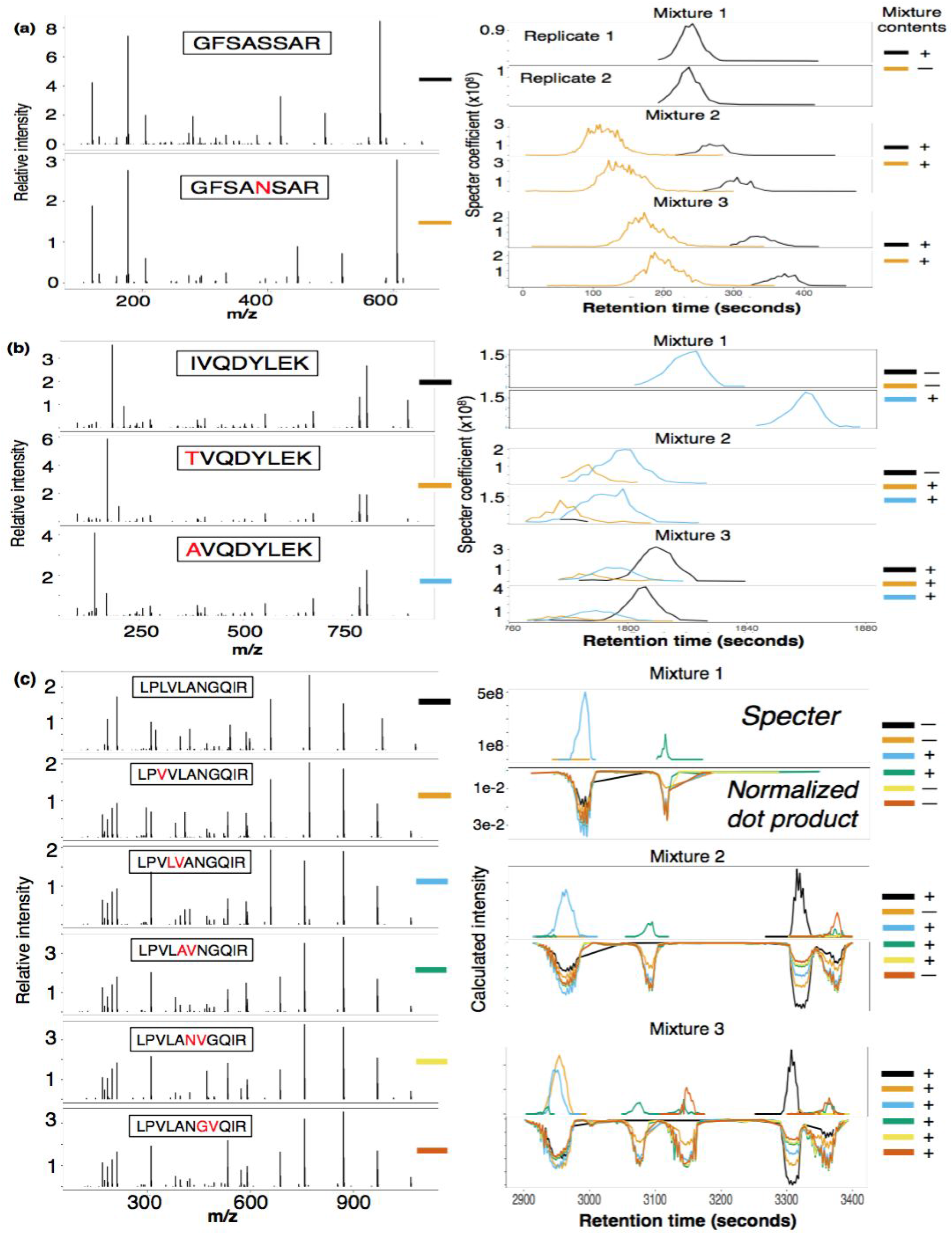
Specter chromatograms of groups of synthetic peptides with highly similar spectra. Members of each of three groups of highly similar peptides (spectra shown on left) were chosen at random to be spiked into both an *E. coli* lysate and neat background in each of twelve DIA runs. Chromatograms for each peptide calculated by Specter in each replicate run of each mixture are shown on right. Actual combinations of peptides in runs are indicated on far right, with each color corresponding to a distinct member of each group (only data from runs in the *E. coli* background are shown). (a) A single amino acid substitution. (b) Two unique substitutions at the N-terminal position, creating identical y-ion series for all family members. (c) A larger family consisting of substitutions and transpositions at various positions in the sequence. Comparison of chromatograms for each peptide in this family as calculated by Specter vs. normalized dot product for each mixture (only data for one replicate per mixture is shown).

Specter is able to distinguish between the synthetic precursors within each family despite their extremely similar library spectra, the cofragmentation of several precursors within each group (indicated by overlapping chromatograms), and the presence of the complex *E. coli* background (Fig. 4; results for the neat background are similar but not shown). Even in cases where coelution is not observed, the fact that library retention time information was not used for the Specter analysis demonstrates its ability to separately identify precursors based purely on features of MS2 spectra. In contrast, the normalized dot product is unable to disambiguate the members of a group of six peptides with extremely similar spectra (Fig. 4c). Specter correctly identified all but one peptide (LPVLANVGQIR) in all runs.

We had the prior expectation that this unidentified peptide would be problematic. To construct the spectral library for this experiment, data were acquired for each peptide in isolation and subjected to DDA-style database search with Spectrum Mill to assign peptide-spectral matches. LPVLANVGQIR was the only one of the eleven synthetic peptides whose fragmentation failed to yield any unique product ions of sufficient intensity for Spectrum Mill to unambiguously distinguish its spectrum from those of the other members of its group. Instead, Spectrum Mill assigned the sequence LPVLA<u>VN</u>GQIR and/or LPVLAN<u>GV</u>QIR to the spectrum derived from LPVLANVGQIR in spite of repeated acquisition. It is very likely that the spurious peaks observed for the peptides LPVLA<u>VN</u>GQIR and LPVLAN<u>GV</u>QIR between 3300 and 3400 seconds are due to LPVLANVGQIR’s ambiguous fragmentation profile (Fig. 4(c)).

### 2.4 Distinguishing positional isomers in phosphoproteomics data from biomedical samples

To illustrate Specter’s ability to distinguish similar precursors in a real-world application, we found examples of positional isomers (peptides with identical amino acid sequences but with post-translational modifications in different positions) with overlapping elution profiles in DIA data that are separately identified and quantified by Specter. This type of analysis is challenging for DDA approaches depending on where fragmentation spectra are sampled during elution, and it has only recently been explored for DIA data^21^. We analyzed 84 DIA runs of a set of phosphopeptide-enriched samples obtained from PC3 prostate cancer cells subjected to a panel of 28 kinase pathway inhibitors in biological triplicate on a Thermo Q-Exactive Plus HF. Analysis was performed using a 12,546-member phosphopeptide library constructed from ten DDA runs of the phosphoproteome of PC3 cells subjected to a subset of the kinase inhibitor perturbations. This library contained 176 sets of unambiguous positional isomers, as determined by Spectrum Mill’s variable modification location score (Methods §4.12).

Specter identified both of the positional isomers GYYS[+80]PYSVSGSGSTAGSR (S4613, in reference the position of the phosphorylated serine on the underlying protein) and GYYSPYSVS[+80]GSGSTAGSR (S4618) in 75 of the 84 runs, and for most of these cases the isomers’ elution profiles overlapped in retention time (Figure 5(a)). We used Specter to disambiguate and quantify the ratio of ion currents from the two positional isomers, which appear to follow disparate patterns of phosphorylation across the perturbations (Figure 5(c)).

**Figure 5:**
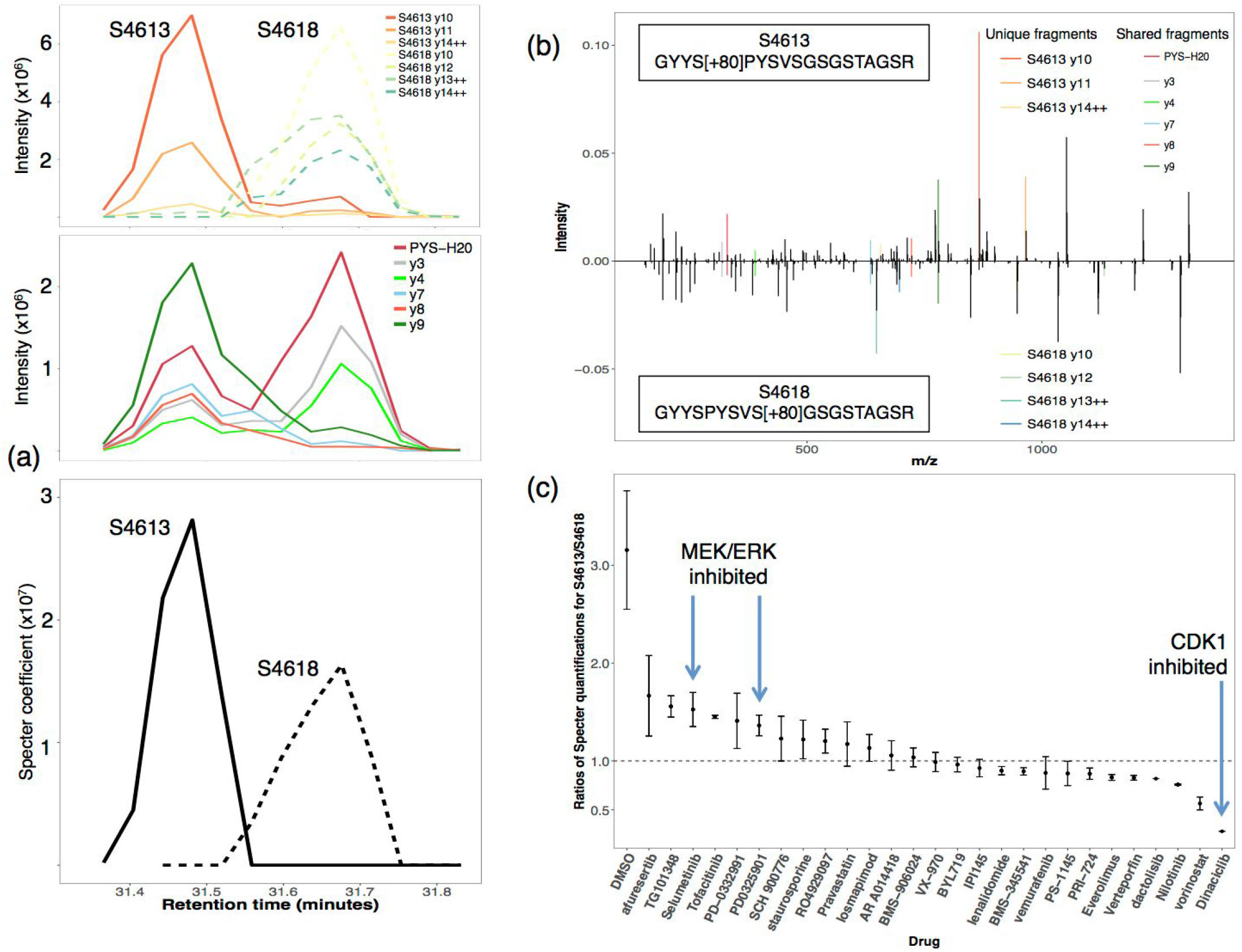
Specter distinguishes close positional isomers with overlapping chromatographic profiles. (a) *Top, first panel:* Raw extracted ion chromatograms for the unique fragments from each of the positional isomers in a DIA run of PC3 cells treated with TG101348. *Top, second panel:* Extracted ion chromatograms for the shared fragments from the isomers. *Bottom:* Specter chromatograms for each of the positional isomers. (b) The spectra of the positional isomers. (c) Mean ratios across replicates of the quantifications of the isomers by Specter (S4613/S4618) across twenty-eight chemical perturbations. Error bars indicate standard error across replicates.

The peptide GYYSPYSVSGSGSTAGSR is a constituent of the cytoskeletal cross-linking protein plectin, located in the last of six highly homologous repeat domains comprising plectin’s C-terminus. The sequence SPYS within this peptide is a known binding motif for CDK1, and it has been shown by site-directed mutagenesis that CDK1 phosphorylates plectin somewhere in repeat domain 6^22^. This phosphorylation was believed to occur at threonine 4,539 based on analysis of CDK1 binding motifs. However, the decreased ratio of S4613/S4618 after treatment with the CDK inhibitor dinaciclib indicates that S4613 (the first serine of the SPYS motif) may be the actual target of phosphorylation by CDK1, or may be cophosphorylated with T4539 given the proximity of these residues in repeat 6.

At the same time, plectin is also known to be a substrate of MAP kinase-interacting serine/threonine-protein kinase 2 (MNK2), which targets a site in repeat 6 distinct from the target site of CDK1^22,23^. Several of the perturbations we analyzed (selumetinib and PD0325901) target the MEK/ERK pathway, part of the MAPK signaling cascade that could regulate the activity of MNK2. Supporting this, ERK inhibitors have been shown to prevent plectin phosphorylation by MNK2^23^. In contrast, the mean S4613/S4618 ratio is lower for the p38 inhibitor losmapimod than for either MEK inhibitor, and p38 inhibitors do not prevent MNK2 phosphorylation of plectin^23^. Taken together, these considerations suggest that serine 4,613 is phosphorylated by CDK1 and not by MNK2, with the converse being true of serine 4,618.

### 2.5 Label-free quantification by DIA with Specter is more reproducible and exhibits a broader dynamic range than DDA

A central pair of premises underlying DIA are that it allows for the detection of analytes with low relative abundance and does not suffer from the inconsistencies of stochastic precursor selection^24^. To test these principles with Specter, an unfractionated *E. coli* lysate was measured with both DDA and DIA strategies, each performed back-to-back in triplicate on the same instrument (Methods §4.9). The DIA runs were analyzed by Specter using a spectral library containing 48,131 precursors obtained from DDA runs of ten fractions of the lysate (the same library as used in §2.2 and §2.3), while analysis of the DDA runs was performed using MaxQuant (Methods §4.11). The MaxQuant “match between runs” option was disabled in order to examine replicate reproducibility directly without *a posteriori* resolution of missing values.

We found that DIA with analysis by Specter is more reproducible than DDA, due to large numbers of precursors which are identified in one replicate DDA run but not another, or which exhibit high variability in their quantifications between runs (Supplementary Figure 4(a)). In contrast, identification and quantification in DIA by Specter are highly reproducible, while the total numbers of peptide and protein identifications (12,204 and 1,190, respectively) are comparable to those obtained with DDA (14,407 and 1,350) in the common precursor range of 389-1015 m/z (we only consider a protein to be identified if at least two of its unique peptides are identified). These observations are quantified by Pearson correlation coefficients (Supp. Fig. 4(b); average r^2^ across DDA replicates = 0.72, average r^2^ across DIA replicates = 0.98). The broader dynamic range of DIA is quantified by the distributions of relative precursor quantifications. The dynamic range of DDA spans roughly four orders of magnitude (∼1.6x10^6^ −2.5x10^10^), while that of DIA with Specter spans more than five (∼3.5x10^5^ −7.4x10^10^).

### 2.6 Comparison to other DIA analysis methods with LFQBench

We used Specter to analyze a publicly available dataset generated for LFQBench^9^, an R package that enables the comparison of label-free quantification results of the five most commonly used DIA analysis tools: OpenSWATH^3^, Skyline^25^, Spectronaut^7^, DIA-Umpire^4^, and PeakView (aka SWATH 2.0). These data were obtained by mixing together the proteomes of three species (human, yeast, and *E. coli)* in two different samples, A and B, at defined ratios and analyzing the mixtures by SWATH^26^ on an AB SCIEX TripleTOF 6600 with 64 variable-width windows. Successful analysis should recapitulate, on average across all identified peptides, the known ratios at which the proteomes were combined (namely 2:1 for yeast, 1:1 for human, and 1:4 for *E. coli).*

We used a spectral library provided by the study’s authors (Methods §4.12) in order to make the most direct comparison possible to the reported results. Specter identified 40,343 of the 44,294 library peptides, corresponding to 4,733 proteins (where we only consider a protein to be identified if at least two of its unique peptides are identified in the same sample). The ranges of the number of identifications by the other library-based tools were 35,517-42,439 peptides and 4,518-4,692 proteins. 677 library peptides were identified by Specter only, the most of the five tools (Figure 6(a)). Figure 6 (b) displays the log-ratios log_2_(A/B) of the most precise peptide quantifications between the two samples (coefficient of variation across replicates < 10%) as reported by Specter and the five other tools as functions of the peptides’ intensities in sample B. The horizontal dotted lines indicate the log-transformed ratios at which the proteomes were mixed in the two samples. We calculated the accuracy of quantification by Specter following the LFQ Bench study methods and compared to the other tools (Supplementary Table C)^9^. Specter is more accurate than all five of the other tools in quantifying the *E. coli* peptides, which have the most extreme expected ratio (1st, 2nd and 3rd tertile mean accuracies of the five non-Specter tools = 0.635, 0.38, and 0.182 vs 0.16, 0.16, and 0.18 for Specter, respectively).

The maximum similarities (normalized dot products) between the unique Specter identifications and all other library spectra are, in a statistically significant fashion, higher overall than these values for the unique Spectronaut identifications (Supplementary Figure 5), with a mean of 0.74 as opposed to 0.64 for Spectronaut (p-value of one-sided Kolmogorov-Smirnov test = 5.24 x 10^−16^). This shows that the 677 unique peptide identifications from Specter complement the identifications from the other tools by including precursors that other methods might deliberately exclude or inaccurately identify due to their similarity to other analytes.

## 3 Discussion

The mixed mass spectra produced by DIA are, in ideal terms, linear combinations of pure spectra. While the constituents of such a combination and their abundances are difficult to measure precisely due to shared fragments, biochemical noise, instrument inaccuracy, and inconsistencies in a peptide’s fragmentation profile across experiments, we have shown that a linear model enables a principled and effective approach to spectral deconvolution. We have shown that Specter integrates the identification of precursors with their quantification, since these are achieved simultaneously through Equation (1) - only precursors with nonzero coefficients can be considered identified, and these coefficients at the same time quantify the intensity of each precursor. This allows for the calculation *in silico* of total ion chromatograms for all library precursors without recourse to individual fragment ion traces, which is an important feature in cases where the user’s spectral library lacks fragment ion annotations.

**Figure 6:**
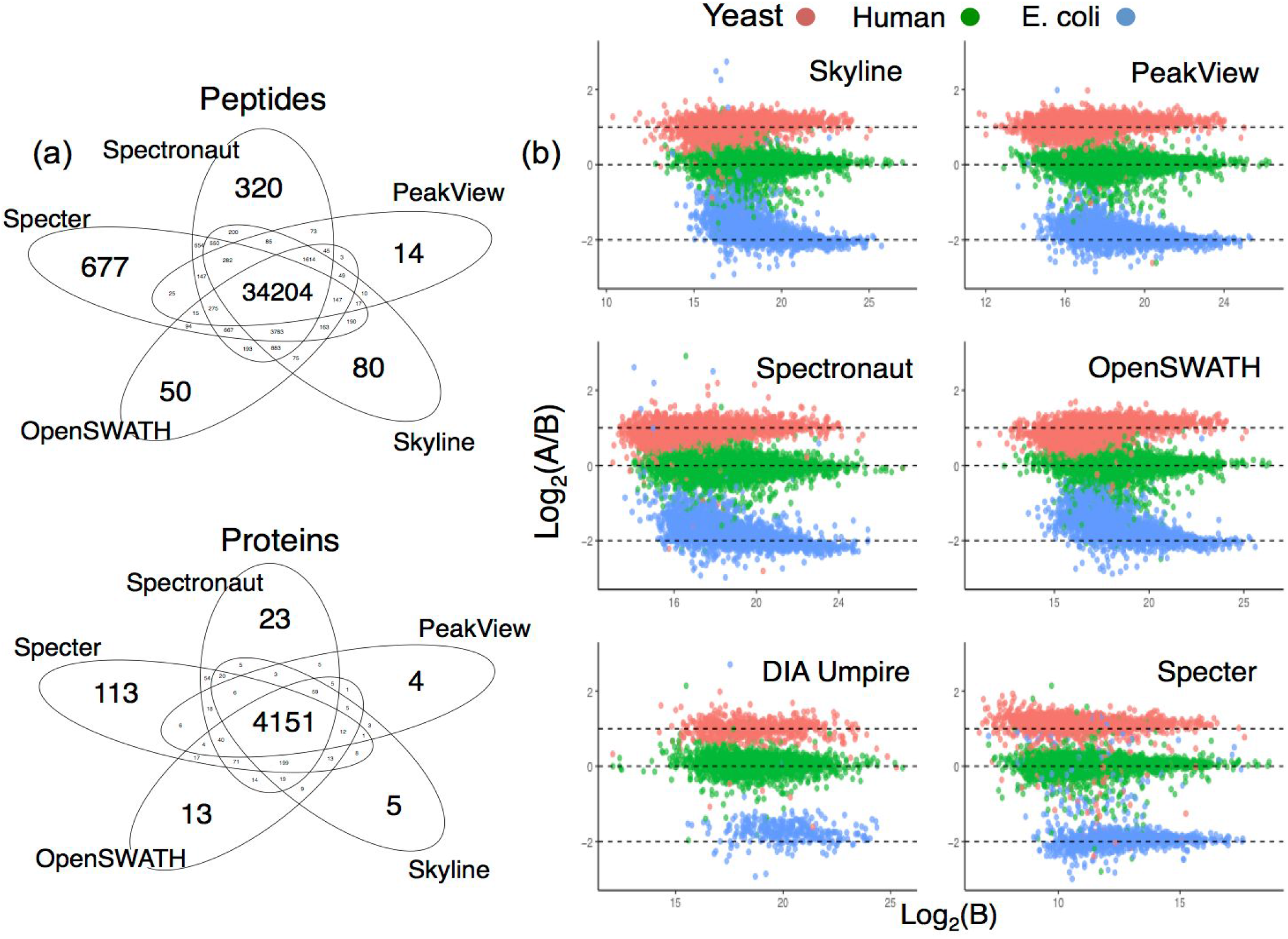
Analysis of a public dataset by Specter and five other software tools. (a) Overlap in peptide and protein identifications from Specter and the four library-based tools. At least two peptides associated to a given protein must be identified in order to consider the protein identified. (b) Log-ratios of high-precision peptide quantifications (CV across replicates < 10%) for the two LFQ Bench samples from all six tools, plotted against intensity in sample B. The expected logarithms of these ratios are indicated by the horizontal dotted lines. Specter quantifications are shown as is, while those for all other tools were scaled to a common range by the authors of the LFQ Bench study.

Specter exhibits a novel ability to separately identify and quantify precursors with highly similar library spectra, even when these precursors are coisolated and cofragmented. As far as we are aware, this type of analysis has not previously been directly shown to be possible for DIA data, despite its potential importance from the perspective of biomedical discovery. In fact, several existing methods include rules to explicitly disallow such cases from consideration by either discarding shared fragments or excluding all but one member of a given group of precursors with similar spectra^3,5,6^.

In terms of practical advantages for working biochemists, our analysis of differential regulation of positional phospho-isomers across a series of perturbations of prostate cancer cells by kinase pathway inhibitors illustrates the kinds of biological insights that may be gained as a result of Specter’s ability to distinguish between similar library analytes while maintaining a low false discovery rate. Furthermore, Specter’s robustness against incompleteness and noise within a spectral library reduces the need for fractionation and time-consuming curation of fragment ions. Given Specter’s use of all features in precursor fragmentation patterns, in general the ideal spectral library for a given DIA experiment should be generated from DDA runs of the samples under consideration using the same instrument.

In the future, we aim to increase Specter’s scope to allow for the characterization of non-library analytes based on correlations between fragment ion elution profiles, in a spirit similar to DIA Umpire^4^. This will expand on the linear model of Equation (1) to explicitly account for the linear contributions of unknown analytes to sequential MS2 spectra while simultaneously identifying and quantifying known library members. Specter is compatible with every instrument type and acquisition scheme, and is available as an open-source software tool (https://github.com/rpeckner-broad/Specter). We also intend to introduce an online interface to which researchers may submit their data for analysis by Specter for users without access to a computing cluster or Apache Spark.

Specter helps to fulfill DIA’s promise to provide numbers of peptide and protein identifications comparable to DDA while exhibiting greater reproducibility across runs and a broader dynamic range. We expect that its high sensitivity and specificity will accelerate the pace of research both into DIA methods generally and into novel applications enabled by the unbiased, reproducible observation and differential quantification of proteins in a broad spectrum of biological contexts.

## 4 Materials and methods

All mass spectrometry proteomics data have been deposited to the ProteomeXchange Consortium via the PRIDE^27^ partner repository with the dataset identifier PXD006722, and can be accessed using the credentials

`Username:`

`Password:` IJfZeE4w

### 4.1 Mass spectrometry data processing

All raw mass spectrometry data files (in either Thermo RAW or AB Sciex WIFF format) were converted to mzML format using ProteoWizard MSConvert version 3.0.6141 with peak picking (centroiding). Spectral libraries are accepted in Skyline’s blib format^25^, which can be constructed from any of the common MS search results file formats^28^.

### 4.2 Python environment and parallelization over MS2 spectra with Apache Spark

Specter is written in Python 2 and runs on Apache Spark, a highly efficient cluster computing framework that enables the parallelization of Specter’s core algorithm over all MS2 spectra acquired in the course of a DIA run. All analyses in this paper were performed using Python 2.7.11 and Spark 1.6.0 on a computing cluster with 48 identical cores (Intel Xeon CPU E5-2697 v2 @ 2.70GHz) and 250 GB RAM. mzML files are exposed to Python using the run.Reader() function from the Python package pymzml v. 0.7.7. Python v. >= 2.7.9 and v < 3 is required to guarantee compatibility with Spark and the packages used by Specter. A Python list is constructed, each of whose elements is a two-column matrix containing the *m/z* coordinates and intensities of the peaks of an acquired MS2 spectrum, and this is converted to a Spark resilient distributed dataset (RDD) via the parallelize() method. The Spark mapPartitions() method is then applied to this RDD to distribute the analysis of the individual MS2 spectra over the computing cluster. The results are returned to the driver node as a list via the collect() method, and subsequently written to a csv file containing the Specter coefficients of each precursor within each MS2 spectrum for downstream processing. A typical DIA experiment file (∼3-5 GB) can be analyzed with a spectral library of ∼20,000 precursors in under thirty minutes by this workflow.

### 4.3 Constructing the reference spectra library matrix

To construct the library spectra matrix *L* i n Equation (1), an instrument mass accuracy parameter δ is required. For the data analyzed in this paper, all of which was acquired on high resolution instruments (Thermo Q exactive Orbitrap or AB Sciex TripleTOF 6600), δ was set to 10 parts per million (ppm) for Orbitrap data and 30 ppm for TripleTOF data. Let *S* be an acquired MS2 spectrum from the DIA run. *S* is analyzed using only a subset of the provided spectral library, since there are physical constraints on the possible presence of a given library member in *S*. The set of library members used to analyze *S* is determined by the following conditions (where *L* denotes a candidate library precursor):

1. The *m/z* ratio of *L* must lie inside the precursor isolation window from which *S* was acquired.
2. At least five of the *m/z* ratios of the peaks of the spectrum of *L* must appear as *m/z* ratios of peaks *S*.
3. If the library includes retention time information, and the library retention times are directly comparable to those in the DIA experiment (as is the case if e.g. the library was generated from DDA runs of the same samples on the same instrument, or both the library and acquired spectra have had their retention times normalized), then the library retention time for *L* must be no more than five minutes greater or less than the time of the scan (this time window can be omitted or adjusted by the user).

While retention time information in the library is optional, it both speeds the analysis by limiting the set of precursors considered for each scan and improves the quality of the results, and so is highly encouraged to be included in cases that the library and DIA spectra are gathered in comparable timeframes.

For each MS2 scan *S*, the *m/z* coordinates of the peaks of the library spectra are then binned with the *m/z* coordinates of the peaks of *S* to obtain a vector of intensities whose length equals the number of peaks of *S* (Supplementary Note 1.1). Each library spectrum is normalized so that its total ion intensity is one, and these normalized spectra are arranged as the columns of a matrix *L* whose number of rows equals the length of *S.*

### 4.4 Finding the optimal combination

Let *S* be an MS2 scan from the DIA experiment, represented as a vector of *n* intensity values. In order to account for peaks of library spectra not matching peaks in S, as described above, we append a zero to the end of this vector so that it has length *n+1.* This extra zero serves to penalize the linear contributions of library spectra that have peaks with significant intensities that aren’t present in *S.* Let *L* be the corresponding matrix of normalized reference spectra constructed above (see also Supplementary Note 1.1). Our aim, as stated in Equation (1), is to find the nonnegative linear combination of the columns of *L* (the normalized library spectra) that best explains *S*, i.e. is closest to it in terms of Euclidean distance. Some peaks of *5* may not be close to any of the peaks of the reference spectra, as determined using the mass accuracy δ, and these may be discarded from the analysis as they don’t affect the determination of the optimal linear combination (Supplementary Note 1.2), i.e. we project the spectrum *5* to the linear span of the library spectra prior to analysis.

With these unnecessary peaks removed, the optimal linear combination of the reference spectra is determined as the solution of the corresponding nonnegative least squares problem, which finds the vector *c* of length *m* (where *m* is the number of spectral library members), all of whose entries are nonnegative, such that the matrix product of *L* with *c* is as close as possible to *5* in the Euclidean norm among all such nonnegative vectors (Supplementary Note 1.2).

### 4.5 Peptide identification from Specter coefficients

From the mathematical formulation above, we see that for every MS2 spectrum *S* acquired in the DIA run, Specter produces a vector c of nonnegative coefficients, each of which is associated to a particular precursor in the spectral library. Each coefficient associated by Specter to a library member in a given MS2 spectrum may be directly interpreted as the sum of the intensities of the fragments produced by that member’s precursor within that spectrum. This is a straightforward consequence of the fact that the library spectra are normalized to have total ion intensity one (Methods §4.3): when such a normalized spectrum *L* is multiplied by a coefficient *c* (meaning that the intensities of all of its peaks are multiplied by this constant), it’s clear that the total ion intensity of the resulting scaled spectrum *cx L* is nothing other than c. Since the aim of Specter is to represent every acquired DIA MS2 spectrum *S* as a linear combination (Equation 1)

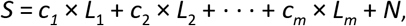

it follows that the total ion intensity of the *i*-th library spectrum *L_i_* in *S* is simply the Specter coefficient *c*¡. Indeed, the multiplication of a library spectrum *L* by a coefficient *c* is the mathematical analogue of the physical fragmentation of *c* molecules of the precursor whose library spectrum is *L*.

Considering all MS2 spectra sequentially, this gives us an *m × r* matrix of coefficients, where *m* is the number of members of the spectral library and *r* is the number of MS2 scans. Each row of this matrix is then a time series describing the “elution profile” of Specter coefficients of a library precursor across the course of the experiment (so that most entries of each row are zero). We consider a library precursor to be identified by Specter if this elution profile contains a peak (local maximum) of at least five consecutive coefficients which are larger than 1 (where coefficients are only considered consecutive if they are calculated relative to sequential MS2 spectra for which the precursor satisfies the conditions of §4.3). This is a physical constraint that recognizes that total ion intensities smaller than 1 can’t possibly correspond to meaningful signal.

As it is possible that peaks such as these may arise by chance, giving rise to false identifications, we employ several chromatographic peak scores in order to rank the quality of our identifications and enable a target-decoy approach for FDR estimation^10^. First, define a peak associated to an identified precursor to be a local maximum within a consecutive series of at least five Specter coefficients larger than one; then define the largest peak to be the peak with the highest summit among all peaks. The four scores associated to the precursor are then 1) the Specter coefficient at the apex of the largest peak, 2) the variance of the coefficients within the largest peak, 3) the skewness of these coefficients, and 4) the kurtosis of these coefficients. Equations for these scores are given in Supplementary Note 1.3. Taken as a set, these four scores measure the extent to which the largest peak within the precursor’s Specter chromatogram resembles an ideal Gaussian elution profile^10^. Rather than enforcing a strict match to a Gaussian, however, we use these scores to develop statistics for confident identifications based on the presumably poor peak shape of false positives (§2.2.1). These four scores are combined into a single score via linear discriminant analysis in order to establish cutoffs to separate target and decoy spectra (see §2.2.1 and §4.7 below).

### 4.6 Peptide quantification from Specter coefficients

Since the output of Specter is affected by experimental noise and the presence of precursors for which library spectra may not exist, filtering of the Specter elution profile of each precursor is essential to obtain accurate quantifications. In order to avoid bias arising from parametric filters, we apply a Kolmogorov-Zurbenko filter with three iterations and windows of width three to smooth the Specter elution profile; this is essentially an iterated moving window average^29^. The quantification of the precursor is then calculated as the area under the largest peak of this filtered profile.

### 4.7 Decoy spectra generation

Our approach to the generation of decoy spectra is developed in the spirit of strategies that generate decoy spectra directly from real library spectra, rather than beginning with random transformations of the sequences of library peptides and subsequently generating decoy spectra based on theoretical fragmentation. The set of decoy spectra used to analyze a given *m/z* window is constructed by first choosing a random subset of all library precursors whose *m/z* ratios do not fall into this window, where this subset is chosen to have the same size as the set of true library spectra for this window. We then construct the decoy spectra for this window by shifting the *m/z* coordinates of all peaks of these non-window spectra by 20 m/z. This method combines the main approaches of Lam et al.^17^ and Cheng et al.^17,18^ In order to avoid distorting the quantifications of non-decoy library members through the influence of decoy spectra in Equation (1), we take a two-pass approach: first, a hybrid target and decoy library is used in order to determine linear discriminant score thresholds (based on the set of scores from §4.5) to achieve a false discovery rate below 1%. Specter is then rerun with the target library only, and only identified library precursors whose linear discriminant scores lie above the determined threshold are retained.

### 4.8 Bovine spike-in experiment

800 ul of soluble *S. cerevisiae* proteins were determined by BCA to be at a concentration of 4 ug/ul. This was placed into a vial of PPS (Expedeon Inc.) to make it about 0.1% PPS and 45 ul of 1 M ABC added to give it pH 8. 20 ul of 500 mM TCEP was added and reduction occurred at 37°C 1200 rpm for an hour. The sample was cooled to room temp and 22 ul 500 mM iodoacetamide added for 30 minutes in the dark. The reduced and alkylated protein solution was added sequentially to three vials (each containing 20 ug Promega trypsin) for a total of 60 ug trypsin. This had argon blown over the top to displace oxygen prior to 37°C 1200 rpm overnight. 60 ul of 10% TFA was used to acidify to pH 2 the next day prior to passing it sequentially over three 30 mg Oasis MCX cartridges (Waters Corporation) for cleaning by dual mode solid phase extraction. Prior to the spike-in experiments, the complex matrix was run four times to condition the liquid chromatography column.

An equimolar six protein digest (Bruker-Michrom) was spiked into the *S. cerevisiae* lysate digest over four orders of magnitude (0amol, 10amol, 30amol, 100amol, 300amol, 1fmol, 3fmol, 10fmol, 30fmol). DIA data were acquired on a Q-Exactive HF (Thermo Fisher Scientific). The maximum fill time for an isolation window was set to 60 milliseconds. MS1 and MS2 scans were acquired with resolving power 30,000 and 60,000 respectively. The AGC target for MS1 scans was set to 1e6 ions and 1e5 ions per isolation window for MS1 and MS2 scans respectively. A total of 20 x 20m/z non-overlapping windows were used to traverse the range from 500-900 *m/z*. 1.2 μg of sample was used per injection. A 75 μm inner diameter fused silica column was packed with 40 cm of reversed-phase C12 Jupiter resin (Phenomenex) and used to separate the sample across a 90-minute linear acetonitrile gradient from 0 to 25% Buffer B. Chromatography was performed using an EASY-nLC II (Thermo Fisher Scientific) system set to a flow rate of 250 nL/min. Buffer A was 2% ACN, 0.1% formic acid, and 97.9% water. Buffer B was 99.9% ACN and 0.1% formic acid.

A spectral library was generated by first searching a DDA run (with the same MS1 parameters as above) of the 30 fmol spike-in sample with Spectrum Mill v. B.06.01.201 using a FASTA containing the six bovine protein sequences as well as the UniProt *S. cerevisae* S288c protein sequences. Results were auto-validated to a FDR of 1%. This yielded a PepXML search results file, which was loaded into Skyline v. 3.6.0.10162 in order to generate a spectral library of 959 yeast and bovine precursors in blib format^30^.

#### 4.8.1 False discovery rate estimation

Of the 959 yeast/bovine library precursors, 757 were identified by Specter in all three replicate DIA runs (78% of the yeast/bovine library), while 381 E. coli peptides were identified in all three replicates (0.8% of the E. coli library). Thus, if a decoy library of the same size as the yeast/bovine library were constructed by choosing 959 of the 48,131 E. coli library spectra uniformly at random, the estimated false discovery rate would be

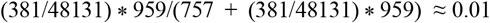

However, when we attempted to actually choose random subsets of 959 precursors from the E. coli library to serve as decoy libraries, we found after twenty such random choices that Specter never assigned a nonzero coefficient to any E. Coli precursor from these smaller subsets (note moreover that a single non-zero coefficient would be inadequate for identification, as it is the time series of such coefficients over the entire DIA run that is assessed to identify precursors, §4.5). This suggests that the 381 false positive E. coli peptides discussed above are symptomatic of the size of the E. coli library, in contrast to the much smaller yeast/bovine library, as this vastly increases the number of degrees of freedom for constructing linear combinations of precursors that most closely match the acquired data.

### 4.9 Synthetic peptide experiment

Synthetic peptides were obtained through New England Peptide Inc. (Gardner, MA), pre-dissolved in 30% acetonitrile (ACN), 0.1% formic acid (FA) and diluted to 100 μM in the same solvent.

To obtain library spectra for the synthetic peptides, each peptide was injected individually at 50 fmol, 200 fmol and 1 pmol on column with wash runs in between. On-line liquid chromatography was performed with an EASY nano-LC 1000 UHPLC (Thermo). Separation was performed on a 20 cm, 75 μm i.d. column, packed in-house with 1.9 micron C18-AQ beads (Dr. Maisch) with a gradient from 2% ACN to 55% ACN over a 20 minute gradient. The data were acquired using a Q-Exactive Plus mass spectrometer (Thermo Fisher Scientific, Waltham, MA) in data-dependent top 12 mode using a resolution of 70,000 for MS1 and 17,500 for the MS2 scans. Dynamic exclusion was disabled to obtain MS2 multiple times for each precursor across the peak. The resulting raw files were searched using Spectrum Mill v. B.06.01.201 with a FASTA containing only the sequences of the synthetic peptides and common contaminants. The best-scoring spectrum for each precursor was then chosen to serve as the precursor’s library spectrum.

DH5*α Escherichia coli* were grown in Luria broth at 37 °C overnight. Cells were pelleted by centrifugation, washed once with cold phosphate buffered saline, flash frozen in liquid nitrogen and stored at −80 °C until processing. To generate *E. coli* lysate digest, the cell pellet was thawed on ice. Once thawed, lysozyme (Sigma) was added to the the pellet and placed on ice with periodic vortexing until viscous. The cells were resuspened in 8 M urea, 50 mM ammonium bicarbonate (ABC) plus protease inhibitors (Roche) and the solution was sonicated with a probe sonicator for 2 min, 3 s on, 2 off, until no longer viscous. After centrifugation at 15,000 x g for 30 min at 4 °C, protein concentration was measured by Bradford assay (BioRad). Disulfide bridges were reduced with 10 mM TCEP (tris(2-carboxyethyl)phosphine, Thermo) and alkylated with 10 mM iodoacetamide (Thermo) for 30 min at room temperature in the dark. The lysate was diluted to 1.5 M urea with 50 mM ABC and digested overnight with a trypsin-to-substrate ratio of 1:100. The digest was desalted using C18 Sep-Pak (Waters). After vacuum centrifugation dried peptides were resuspended to 1 mg/mL in 30% ACN/0.1% FA and stored at −80 °C.

To generate library spectra of the *E. coli* digest, peptides were fractionated using a Stage tip^31^ packed with sulphonated poly (styrenedivinylbenzene) resin (SDB-RPS, EMPORE). 100 μg of digest was fractionated starting from 20 mM ammonium hydroxide, pH 10, increasing the percentage of ACN at steps of 5, 10, 12.5, 15, 17.5 20, 25, 30, 35 and 50%. Assuming equal mass distribution, 1 μg of fractionated digest was analysed by LC-MS2. Data were acquired on the same instruments as above with changes to the LC gradient and the data acquisition. Peptides were separated on-line with an 85 min gradient from 6% 0.1% FA (buffer A) to 30% 0.1% FA, 90% ACN (buffer B), followed by an increase to 60% buffer B over nine minutes. The mass spectrometer was set to acquire data at a resolution of 70,000 and an AGC setting of 3e6 for MS1. MS2 resolution was 17.5K, AGC 5e4, and maximum inject time of 100 ms. The top 12 ions identified within the precursor scan of 300-2000 *m/z* of at least doubly charged were selected for HCD at a normalized collision energy (NCE) of 25. Raw files were searched by SpectrumMill v. B.06.01.201 using the NCBI E. coli K12 DH10B FASTA sequence database (a FASTA for DH5α was unavailable) and auto-validated to a false discovery rate of 1%. The results were exported as a PepXML summary file, which was imported into Skyline v. 3.6.0.10162 to generate a spectral library consisting of 48,131 precursors. This spectral library was exported in blib format^25,28^ for later use by Specter.

For spike-in experiments, synthetic peptide mixtures were constructed according to Table A below. 20 injections worth of peptide mixtures for each of the zero, single, or double drop out samples were created. The pools were then equally divided either as synthetic peptides alone, or into the same *E. coli* digest described above. In either case, roughly equal amounts of synthetic peptide material was loaded on column, regardless of whether or not *E coli* background was present. For runs containing lysate background approximately 1.5 μg of *E. coli* digest was loaded on column. Data-dependent acquisition was performed on a Q-Exactive Plus HF mass spectrometer, where MS1 scans were measured at a resolution of 60,000 and an AGC setting of 3e6 and maximum injection time of 20 ms. MS2 scans on the top 15 peaks doubly charged and above were acquired at a resolution of 15,000, AGC target of 5e4 and maximum inject time of 50 ms. Isolation widths were set to 1.7 Th with a 0.3 Th offset. NCE was set to 28 and dynamic exclusion was set to 15 s.

Data-independent acquisition (DIA) data were acquired with MS1 parameters as above (range: 300-1200 m/z) and then using 22 Th wide windows for MS2 with a default charge state of 4 at a resolution of 30,000. AGC target was set to 1e6, maximum inject time was set to 50 ms, and a loop count of 27. NCE was set to 27. A total of 56 x 22 Th DIA windows were used to traverse *m/z* range from 400-1000, wherein the range is actually traversed twice but the windows are offset by 50%. The window centers can be found in Table B (SI). LC and nanospray parameters were identical to those described in Abelin et. al^32^.

**Table A:** Design for the experiment of §2.3. A dot indicates that the peptide was spiked into the indicated mixture.

| Peptide | Mixture 1 | Mixture 2 | Mixture 3 |
| --- | --- | --- | --- |
| GFSASSAR | • | • | • |
| GFSANSAR |  | • | • |
| IVQDYLEK |  |  | • |
| TVQDYLEK |  | • | • |
| AVQDYLEK | • | • | • |
| LPLVLANGQIR |  | • | • |
| LPVVLANGQIR |  |  | • |
| LPVLVANGQIR | • | • | • |
| LPVLAVNGQIR | • | • | • |
| LPVLANVGQIR |  | • | • |
| LPVLANGVQIR |  |  | • |

### 4.10 Positional isomers in drug-perturbed cellular systems

Sample preparation and experimental procedures were identical to those described in Abelin et. al.^32^, and DIA runs were performed with the same parameters used in the synthetic peptide experiment described above in §4.9. Drug treatments and concentrations are shown in Table B below. Ten randomly chosen phosphoenriched samples of PC3 cells treated with the perturbations highlighted in red in Table B were measured by DDA (acquired with the same settings as the DDA runs described in §4.9). Results were searched using Spectrum Mill as above with a FASTA containing the 2014 UniProt Human proteome and 150 common contaminants. Phosphorylation of serine, threonine and tyrosine were set as variable modifications. The resulting pepXML files were imported into Skyline to construct a redundant blib containing multiple PSMs for each precursor. Search results were further filtered by variable modification location score so that only spectra for which phosphosites could be unambiguously localized were retained, and the highest scoring spectra for each precursor were extracted from the redundant blib to obtain a nonredundant spectral library consisting of only confidently localized phosphopeptides. Specter was applied to the 84 DIA runs using this spectral library and a 10 p.p.m. mas accuracy.

**Table B:**
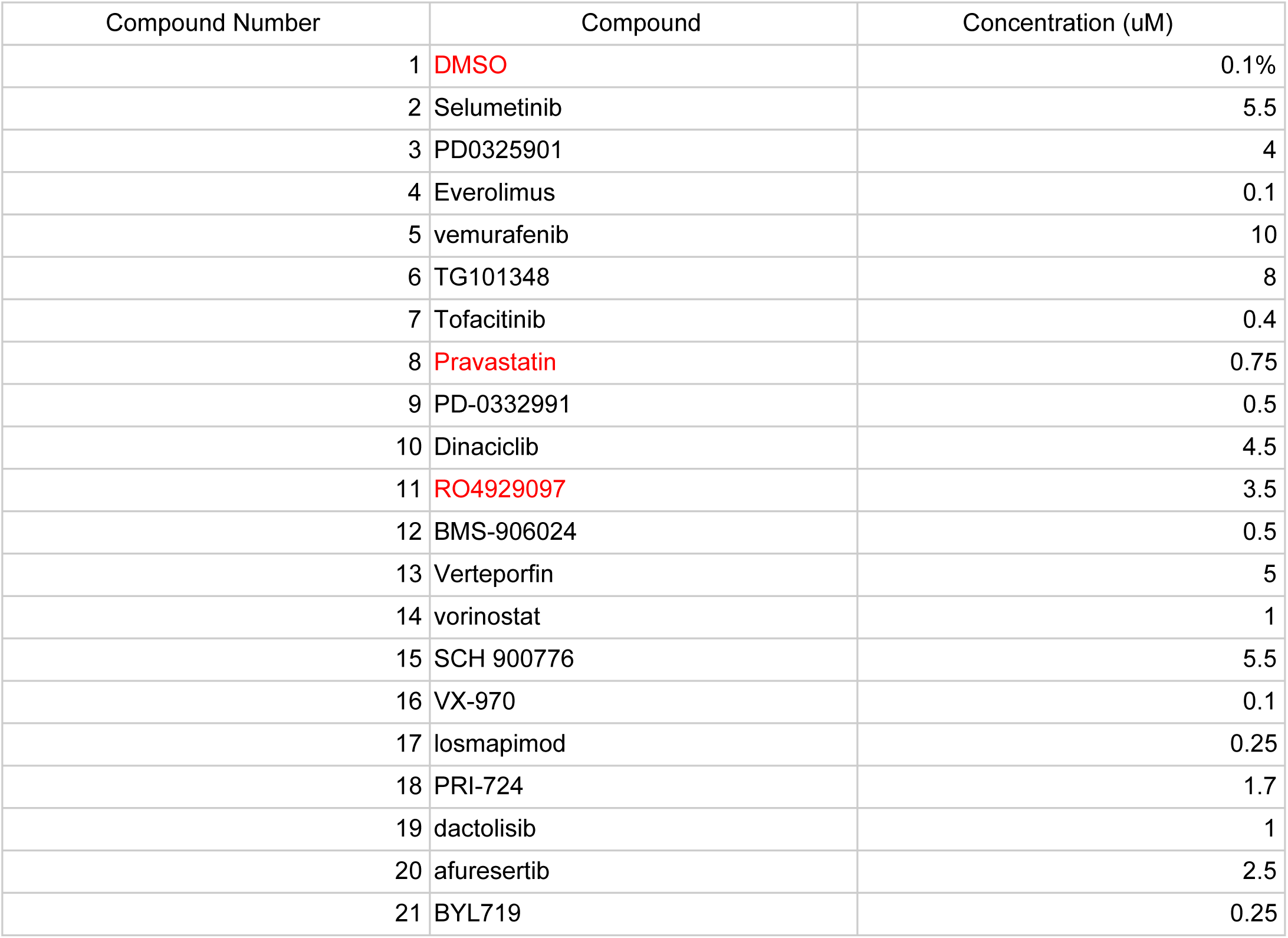
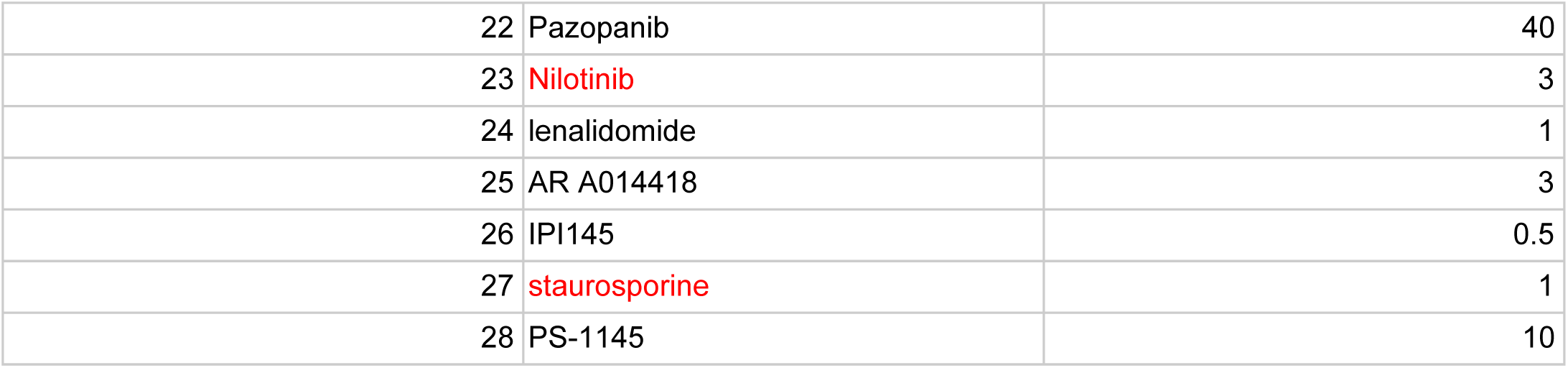
Drug treatments and concentrations for phosphoenriched PC3 samples.

### 4.11 *E. coli* DDA analysis with MaxQuant

RAW files obtained from triplicate DDA runs of the unfractionated *E. coli* digest described above were imported into MaxQuant v. 1.5.5.1. These were searched using the same FASTA as used by Spectrum Mill above with all default settings for Orbitrap instruments *(Match between runs* was disabled in order to assess replicate reproducibility). Results were analyzed using the evidence.txt output table. Only precursors with posterior error probability < 0.01 were retained, and the top scoring MS2 spectrum for each of these precursors within each replicate was used to determine precursor quantifications within each run. Protein identifications (based on all three replicates) were determined from the proteinGroups.txt output table.

### 4.12 LFQ Bench dataset

All data were downloaded from the ProteomeXchange (data set identifier PXD002952). Raw WIFF files from the HYE124 dataset (with 64 variable width windows on a Triple TOF 6600) were converted to mzML using ProteoWizard as described above. We used a spectral library provided by the study’s authors (ecolihumanyeast_concat_mayu_IRR_cons_openswath_64w_var_curated.csv) which consisted of precursors with annotated fragment ions in CSV format compatible with OpenSWATH. The mass accuracy parameter δ was set to 30 ppm.

For comparisons to other analyses, only the results from the first iteration of the LFQBench study were used. This was based on the consideration of the optimizations and open discussion among software developers that took place for the second iteration, in which we did not participate.

## ACKNOWLEDGEMENTS

This work was supported by NIH Grant U54 HG008097 to JDJ. The authors would like to thank Stefan Tenzer and Ludovic Gillet for providing details of the datasets for the LFQ  Bench study.

